# Pan-genome-wide analysis of *Pantoea ananatis* identified genes linked to pathogenicity in onion

**DOI:** 10.1101/2020.07.24.219337

**Authors:** Gaurav Agarwal, Divya Choudhary, Shaun P. Stice, Brendon K. Myers, Ronald D. Gitaitis, Stephanus N. Venter, Brian H. Kvitko, Bhabesh Dutta

## Abstract

*Pantoea ananatis* is a member of a *Pantoea* spp. complex that causes center rot of onion, which significantly affects onion yield and quality. This pathogen does not have typical virulence factors like type II or type III secretion systems but appears to require a biosynthetic gene-cluster, HiVir/PASVIL (located chromosomally), for a phosphonate secondary metabolite, and the onion-virulence regions, OVR (localized on a megaplasmid), for onion pathogenicity and virulence, respectively. We conducted a deep pan-genome-wide association study (pan-GWAS) to predict additional genes associated with pathogenicity in *P. ananatis* using a panel of diverse strains (*n* = 81). We utilized a red-onion scale necrosis assay as an indicator of pathogenicity. Based on this assay, we differentiated pathogenic (*n* = 51)- vs. non-pathogenic (*n* = 30)-strains phenotypically. Pan-GWAS revealed a large core genome of 3,153 genes and a flexible accessory genome of ≤5,065 genes. Phylogenomic analysis using pan-GWAS and presence and absence variants (PAVs) distinguished red-scale necrosis inducing (pathogenic) strains from non-scale necrosis inducing (non-pathogenic) strains of *P. ananatis*. The pan-GWAS also predicted 42 genes, including 14 from the previously identified HiVir/PASVIL cluster associated with pathogenicity, and 28 novel genes that were not previously associated with pathogenicity in onion. Of the 28 novel genes identified, eight have annotated functions of site-specific tyrosine kinase, N-acetylmuramoyl-L-alanine amidase, TraR/DksA family transcriptional regulator, and HTH-type transcriptional regulator. The remaining 20 genes are currently hypothetical. This is the first report of using pan-GWAS on *P. ananatis* for the prediction of novel genes contributing to pathogenicity in onion, which will be utilized for further functional analyses. Pan-genomic differences (using PAVs) differentiated onion pathogenic from non-pathogenic strains of *P. ananatis*, which has been difficult to achieve using single or multiple gene-based phylogenetic analyses. The pan-genome analysis also allowed us to evaluate the presence and absence of HiVir/PASVIL genes and 11 megaplasmid-borne OVR-A genes regarded as the ‘*alt*’ cluster that aid in *P. ananatis* colonization in onion bulbs. We concluded that HiVir/PASVIL genes are associated with pathogenic *P. ananatis* strains and the *alt* gene cluster alone is not responsible for pathogenicity on onion. The pan-genome also provides clear evidence of constantly evolving accessory genes in *P. ananatis* that may contribute to host-range expansion and niche-adaptation.

**Author summary:** *Pantoea ananatis* is a major bacterial pathogen that causes center rot of onion and diseases of a number of other plant species. In order to understand the genome architecture and identify genes responsible for pathogenicity in onion, a pan-genome analysis was performed. We used 81 strains of *P. ananatis* collected over 20 years from different regions of the state of Georgia, USA. The pan-genome study identified a core genome with a conserved set of genes and an accessory genome that displayed variation among strains. We conducted pan-GWAS (pan-genome-wide association study) using presence and absence variants (PAVs) in the genomes and associated onion-pathogenic phenotypes based on a red-onion scale necrosis assay. The study resulted in identification of genes, including a cluster of chromosomal HiVir/PASVIL genes, that are significantly associated with the onion pathogenic phenotype. In addition, we identified 28 novel genes, a majority of which (*n* = 20) have hypothetical functions. We concluded and further substantiated earlier findings that a cluster of genes is responsible for pathogenicity on onion. The pan-genome analysis also allowed us to evaluate the presence and absence of HiVir/PASVIL genes and 11 megaplasmid-borne OVR-A genes regarded as the ‘*alt*’ cluster that aid in bacterial colonization of onion bulbs by *P. ananatis* strains. We concluded that HiVir/PASVIL genes are associated with onion-pathogenic strains, and the *alt* gene cluster alone is not responsible for pathogenicity on onion. This study also provides potential evidence of constantly evolving accessory genes in *P. ananatis* which may help in host range expansion and adaptation to diverse niches.

## Introduction

The genus *Pantoea* currently has 27 recognized species; five of which are known to cause disease-associated losses in several crops [1-5]. Three species of *Pantoea*, namely, *P. ananatis, P. agglomerans* and *P. stewartii* subsp. *indologenes* are responsible for more than 80% of the reported cases of disease in onions [6]. *P. ananatis*, a Gram negative and facultative anaerobic bacterium that belongs to the Erwiniaceae (previously assigned to Enterobacteriaceae) is part of a *Pantoea* spp. complex (also includes *P. allii, P. agglomerans* and *P. stewartii* subs. *indologenes*), which causes center rot of onion [7-9]. Foliar symptoms primarily appear as white streaks and water-soaked lesions, and more advanced infections result in complete collapse of foliar tissues, discoloration and softening of specific scale layers in the bulb. Under favorable conditions, center rot can result in 100% losses in field. In Georgia, late maturing varieties are more susceptible to center rot than early maturing varieties [10-12]. Out of the four species in the *Pantoea* spp. complex, *P. ananatis* has been associated predominantly with center rot, in Georgia [13]. *P. ananatis* has been identified in other onion-growing regions of the United States, including Colorado [14], Michigan [15], New York [16], and Pennsylvania [17]. The bacterium can be found as an epiphyte on crop and weed plants [7] or as an endophyte in maize kernels and rice seeds [18, 19]. Apart from its epiphytic and endophytic niche in crops and weeds, *P. ananatis* can be disseminated through infected onion seed and contaminated insect vectors (thrips) to onion crops [13, 20].

*Pantoea ananatis*, unlike many other phytopathogenic bacteria, lacks genes that code for type II, III and IV protein secretory systems that are associated with pathogenicity and virulence [5]. Recent studies utilized whole genome/small RNA sequencing that aided in identifying some virulence factors associated with *P. ananatis* in onion. These virulence factors are flagellar and pilin motility factors [21] and a global virulence regulator (Hfq, an RNA chaperone) associated with quorum sensing and biofilm production [22]. However, these genetic factors are present in both onion pathogenic and non-pathogenic strains. Recently, a proposed biosynthetic gene-cluster, HiVir/PASVIL (High Virulence also known as PASVIL; *Pantoea ananatis* specific virulence locus) that encodes a proposed phosphonate or phosphinate secondary metabolite cluster located on the chromosome, has been demonstrated to be associated with onion foliar and bulb necrosis [23, 24]. In addition, Stice et al. [25] showed that a megaplasmid-borne onion virulence region (OVR) in *P. ananatis* is correlated with onion virulence. In a recent study, Stice et al. [26] showed that the OVRA cluster contains 11 genes that are critical for colonizing necrotized bulb tissue. This gene cluster was described as the ‘*alt’* cluster that imparts bacterial tolerance to the thiosulfinate ‘allicin’ in onion bulbs.

Whole genome sequencing is a powerful approach used to identify genomic/genetic variants in the genomes of organisms. These genomic variants can be integrated with the unique phenotypic characteristics of an organism using computational methods that may help in identifying genetic determinants that govern the variants. Such computational association methods have been utilized in research of humans [27-29], domesticated plants [30, 31], animals [32], and bacteria [33-35]. This approach is widely used as a genome wide association study (GWAS) that involves genomic characteristics such as single nucleotide polymorphisms (SNPs), insertion and deletions (InDels), copy number variants (CNVs) or the presence and absence variants (PAVs). GWAS studies aim to associate these variants with unique phenotypic characteristics of an organism. In this study, we deployed a pan-GWAS to identify PAVs in *P. ananatis* strains (*n* = 81) associated with onion scale necrosis. Red-onion scale necrosis is a predictor of pathogenicity in onion; hence, we utilized this trait for high-throughput screening to differentiate pathogenic from non-pathogenic strains. We also utilized PAVs to differentiate pathogenic and non-pathogenic strains of *P. ananatis*.

## Results

### The *P. ananatis* genome and pan-genome architecture

A total of 675.6 million raw reads were generated and, after stringent quality filtering, 657.6 million high-quality reads were obtained (S1 Table). Overall, more than 97% of the read data were retained after quality filtering, amounting to 98.6 Gb of data (S1 Fig). These quality-filtered reads were used to construct draft assemblies. We conducted pan-genome analyses on the draft genomes of the 81 selected *P. ananatis* strains collected from onions, weeds and thrips from different regions of Georgia, USA (Table 1). The size of the draft assemblies ranged from 4.7 to 5.8 Mb and the GC content ranged from 52.6 to 53.6%, with strains PANS 99-31 and PNA 18-3S showing the maximum and minimum GC content, respectively. The strains PNA 02-12 and PNA 98-11 had minimum and maximum assembly sizes of 4.7 and 5.8 Mb, respectively (Table 1). In all the assemblies, the total number of contigs ranged from 67 to 2,333, protein-coding sequences ranged from 4,300 to 5,110, and the number of mRNAs (including tRNA, rRNA and tmRNA) ranged from 4,373 to 5,234 (Table 1).

The full spectrum of the pan-genome contained 14,452 clusters of protein coding genes. Among these, 3,153 (present in all 81 strains) were core, 3,799 (present in >=76 genomes) were soft-core, the largest group of 6,808 (<=2 genomes) were cloud and the remaining 3,845 genes belonged to shell gene cluster (Fig 1a and 1b). Each genome had a conserved set of 3,153 core genes and a variable number of accessory genes, which included the soft-core, shell and cloud genes (Fig 1b). Cloud genes are considered to be unique genes contributed by each strain of *P. ananatis* or the class of gene cluster that is next to the most populated non-core cluster of genes. In this study, we found 6,808 novel gene clusters in two or less genomes (Fig 1c). Details of the number of core and accessory genes contributed by each strain are listed in S2 Table.

**Figure 1.**
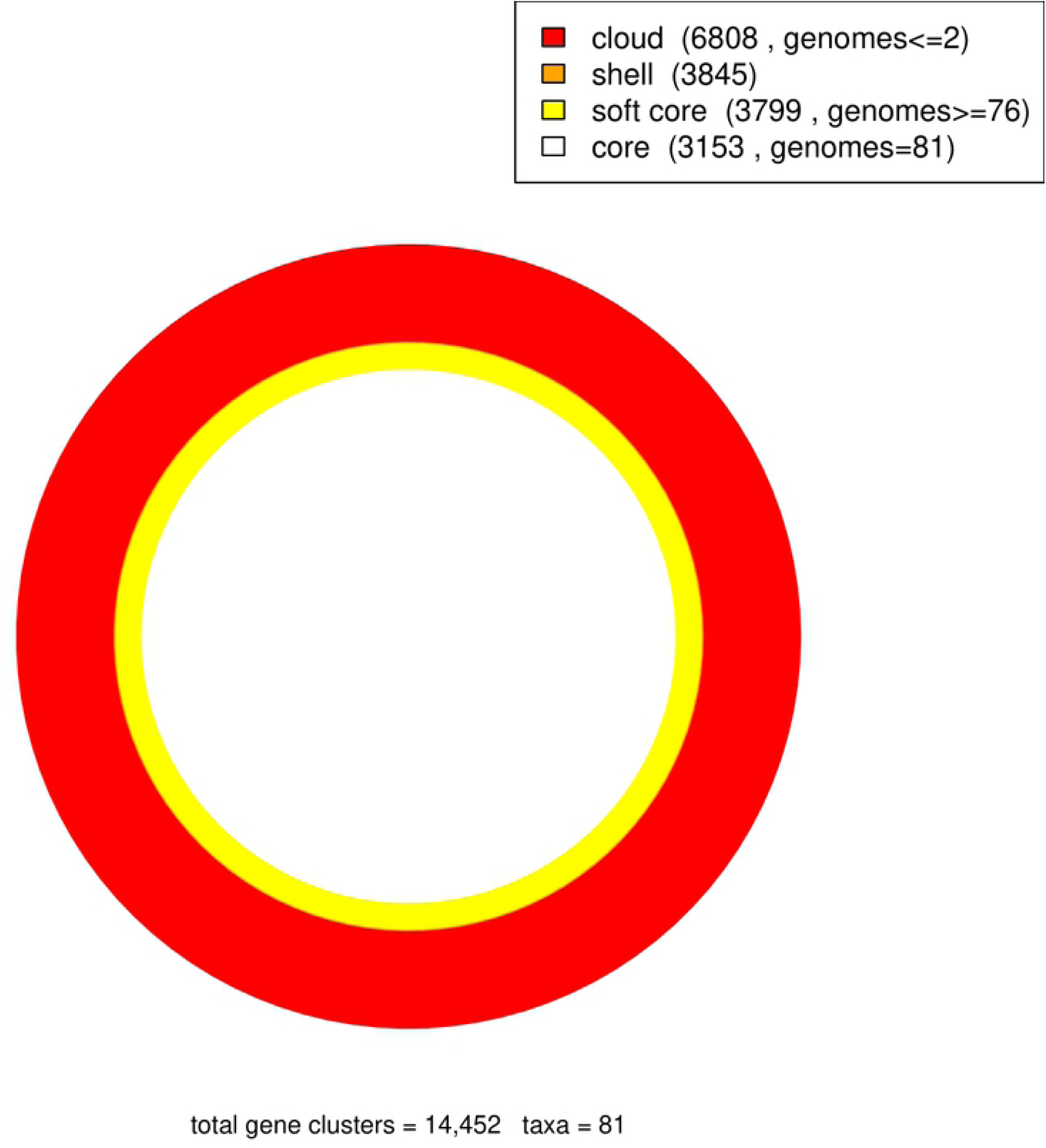
Pan-genome analysis of 81 *Pantoea ananatis* genomes. (A) Area plot of the pan-genome matrix showing the number of genes contributed by core, soft core, shell and cloud genomes. Soft core compartment includes also the core. (B) Genes contributed to pan-genome by individual genomes. (C) Distribution of cluster sizes as a function of the number of genomes they contain showing the partition of OMCL pan-genomic matrix into shell, cloud, soft-core and core compartments.

The Average Nucleotide Identity (ANI) and Average Amino Acid Identity (AAI) are widely practiced genome-based characteristics for understanding the genome architecture of strains within a species. We, therefore, estimated ANI and AAI-based all-vs-all matrices and constructed clustered tree-based heat-maps. Overall, the ANI and AAI among 81 *P. ananatis* strains varied from 99.0 to 99.9% and 96.0 to 99.8%, respectively, which confirmed that all strains used in this study belong to the species *P. ananatis* (Fig 2a and 2b).

**Figure 2.**
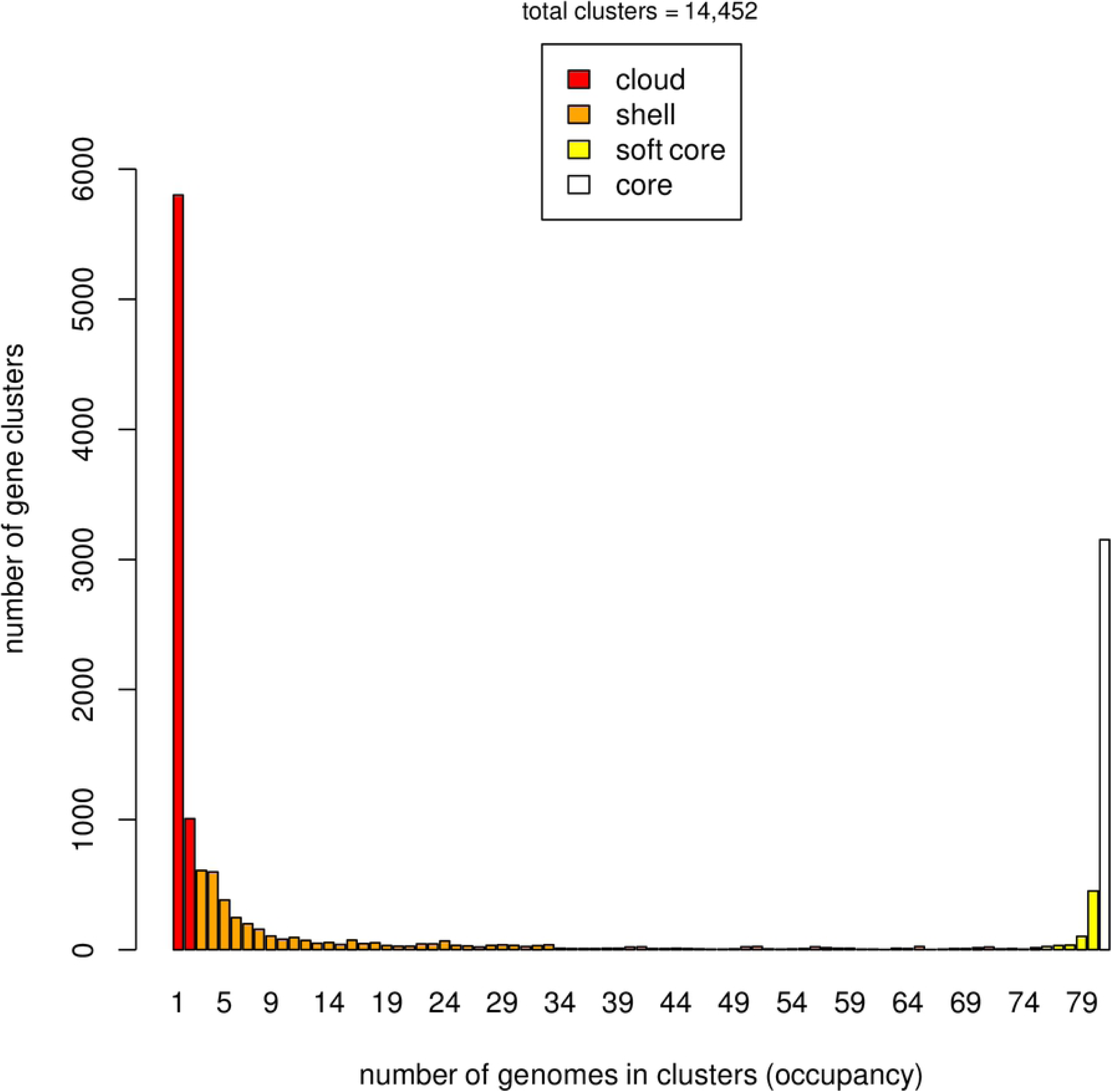
Average nucleotide identities (ANI) of the coding DNA sequences and average amino acid identities (AAI) of the protein coding genes of 81 *Pantoea ananatis*. (A) Heatmap was generated using the identity matrix calculated by get_homologues.pl using the BLAST scores representing the degree of similarity of the genomes based on ANI. (B) Heatmap showing the degree of similarity of the genomes based on the AAI calculated using BLASTP scores implemented in get_homologues.pl. Heatmaps were derived from the ANI and AAI matrix based on the pan-genome matrices. High similarity is represented by lighter color (light yellow to white) and low similarity is represented by dark orange to light orange shade.

### Phenotyping

One hundred percent of the strains used were identified as *P. ananatis* based on PCR assays and biochemical assays, prior to phenotyping. *P. ananatis* strains were classified as pathogenic and non-pathogenic based on the ability to necrotize red onion-scales and clear the anthocyanin pigment. Strains that caused red onion scale necrosis were classified as pathogenic (+) and those that did not were identified as non-pathogenic (-) (Table 1). Based on the red-scale necrosis assay, 62.9% (*n* = 51) and 37.1% (*n* = 30) of the strains were identified as pathogenic and non-pathogenic, respectively. Among the strains that were isolated only from onions (regarded as PNA), 68% (36 of 53 onion strains) were able to cause red-onion scale necrosis (Table 1). In contrast, among the strains (*n* = 28) that were isolated from weeds and thrips (regarded as PANS), 28.5% (8 of 28) and 25% (7 of 28) were able to cause necrosis on the red-onion scale, respectively. A majority of strains (13 of 28) from non-onion sources [weeds: 28.5% (8 of 28) and thrips: 17.8% (5 of 28)] were classified as non-pathogenic as they were not able to cause red-onion scale necrosis (Table 1).

### *P. ananatis* has an open pan-genome

The pan-genome architecture of the 81 *P. ananatis* genomes analyzed is characterized in Fig 3. We used the exponential decay model of Willenbrock [36] that fitted the core gene clusters generated using the OMCL algorithm, which predicted a theoretical core genome of 2,935 genes. In addition, the core genome was not continuous because of the draft assemblies (not the finished sequence assembly) used in this study (Fig 3a). In order to check the openness/closeness of the pan-genome, the theoretical estimation of pan-genome size was carried out using an exponential model of Tettelin et al. (2005) [37], which was fitted to the OMCL accessory gene clusters. The pan-genome samples appeared to converge to linear growth with >10,000 genes, with ∼50 new genes being added on average to the pan-genome with each new *P. ananatis* genome sequenced (Fig 3b).

**Figure 3.**
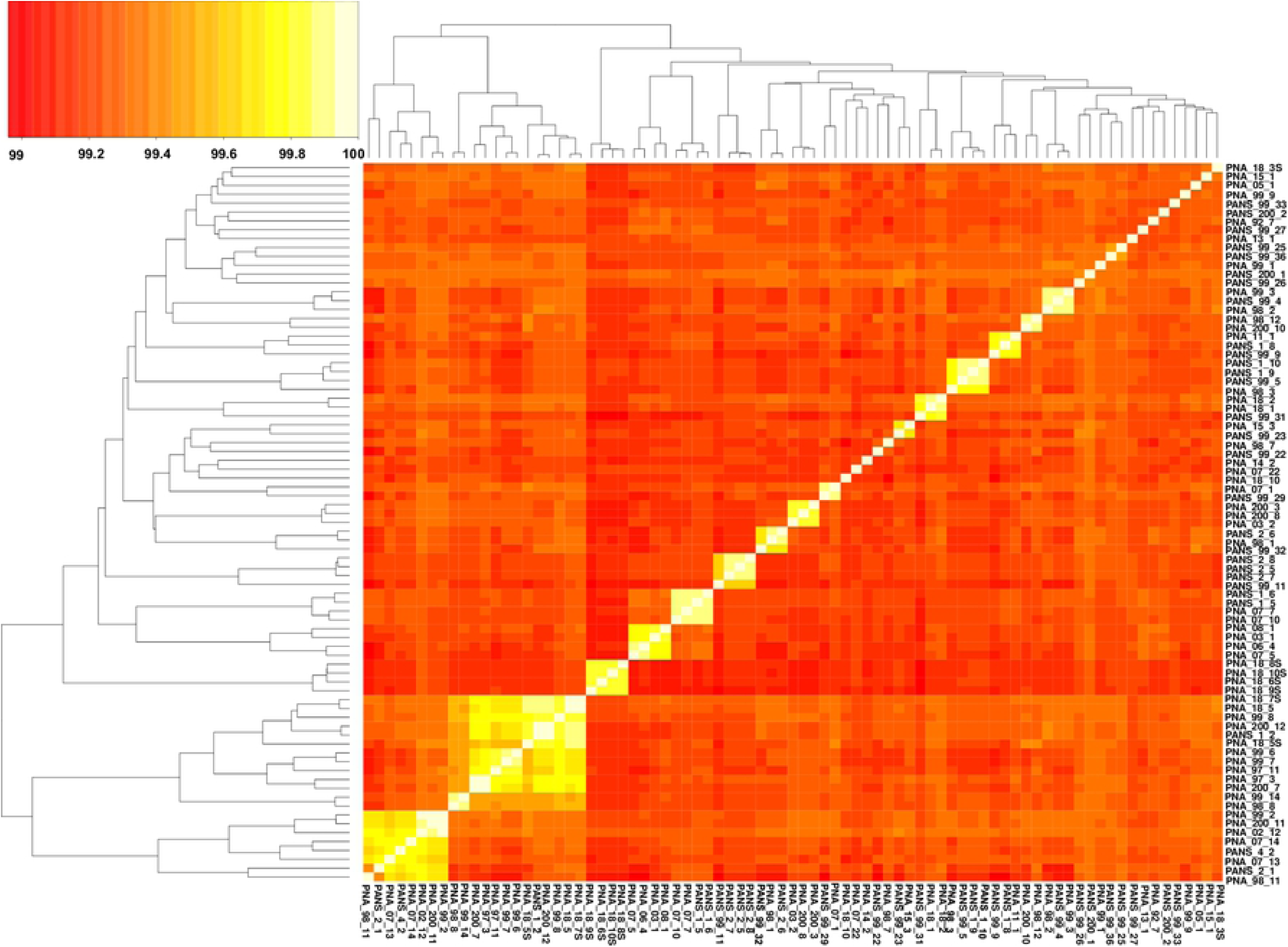
Theoretical estimation of the core and pan-genome sizes based on the exponential decay model. (A) Estimation of core genome size based on Willenbrock model fit to OMCL clusters. (B) Estimation of pan-genome size based on Tettelin model fit to OMCL clusters.

### Analysis of *P. ananatis* strains revealed pathogenic and non-pathogenic differentiation

A pan-genome (including both core and accessory) based dendrogram utilizing PAV genes was constructed to understand the relationship among strains of *P. ananatis* (Fig 4). We used 51 onion pathogenic and 30 non-pathogenic strains as identified using the red-scale necrosis assay stated above. Strains were distributed in four clusters in the pan-genome tree. The first cluster contained 12 pathogenic strains (red) and one non-pathogenic strain (green), the second cluster (II) contained 22 non-pathogenic strains, the third cluster (III) contained 39 pathogenic strains and the fourth cluster (IV) contained seven non-pathogenic strains (Fig 4). The strain PNA_98_11 (non-pathogenic) was the only outlier that clustered with pathogenic strains in the first cluster but otherwise the dendrogram resolved the pathogenic and non-pathogenic strains. We also observed that cluster I of the pathogenic strains contained nine PANS (strains from non-onion sources) and three PNA (strains from onion) strains out of the total 12 strains (Fig 4). However, cluster III of the pathogenic strains contained only six PANS and 33 PNA strains of the total 39 strains. In addition, geographical location, year of isolation, and source of isolation did not show any trend in relation to clustering (Fig 4).

**Figure 4.**
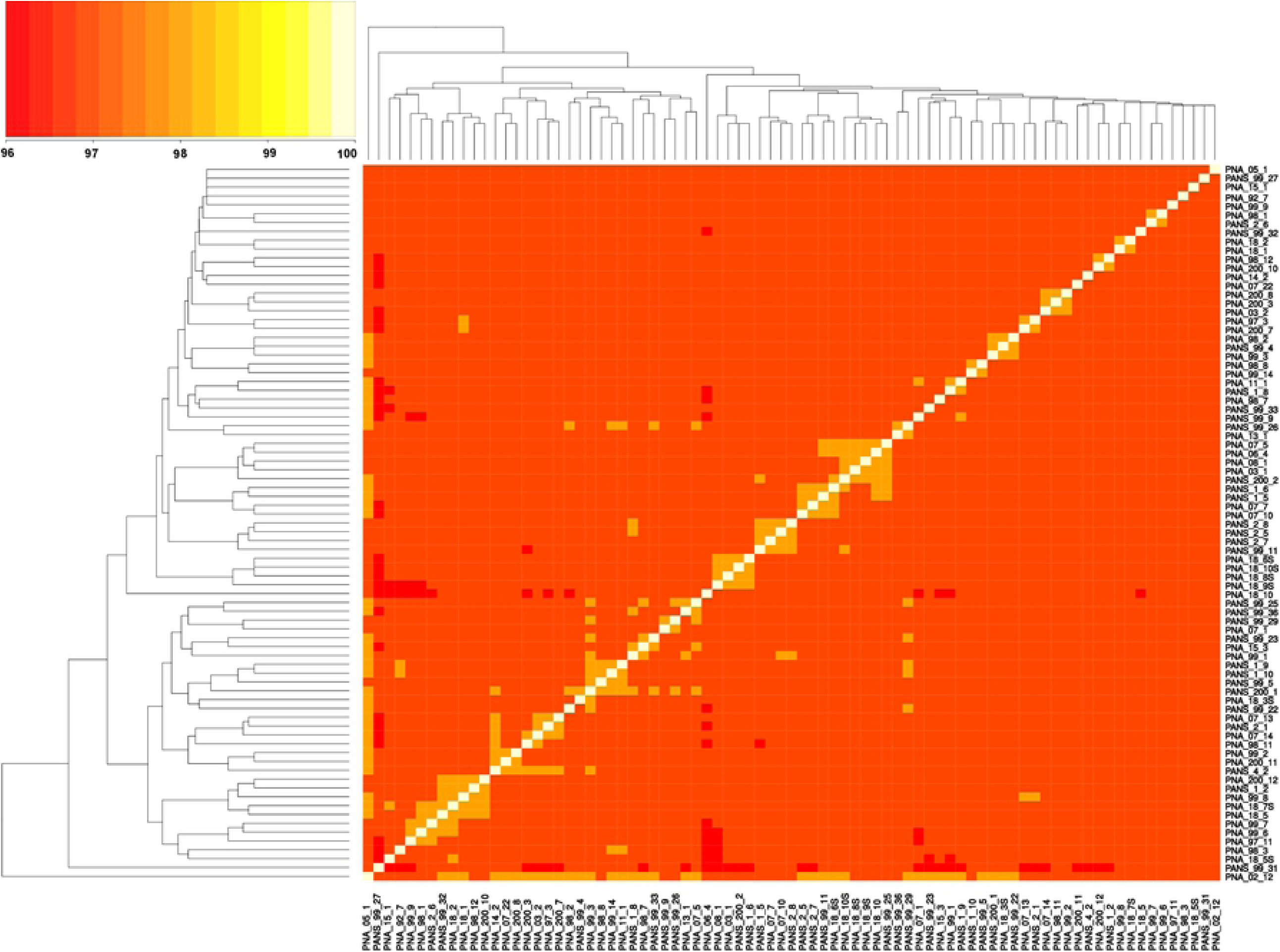
Dendrogram of 81 pathogenic and non-pathogenic strains of *Pantoea ananatis* based on the presence and absence of genes. Strains highlighted in green are non-pathogenic and ones in red are pathogenic.

We also constructed dendrograms using only the core, shell and cloud genes individually to compare the resolution of pathogenic- and non-pathogenic strains for each with that of the entire pan-genome PAVs. Conserved core genes clustered pathogenic- and non-pathogenic strains. Broadly, two clusters were obtained with one cluster comprised of a majority of the non-pathogenic strains [28] and nine pathogenic strains, and the other cluster comprised of 42 pathogenic- and two non-pathogenic strains (S2 Fig). Shell PAVs constituted two broad clusters; one with 47 pathogenic- and 13 non-pathogenic strains, and another with 17 non-pathogenic and four pathogenic strains (S3 Fig). Cloud PAVs showed a mixed pattern of clustering of pathogenic- and non-pathogenic strains (S4 Fig).

### Pan-genome-wide association study

This study identified a conserved set of core and accessory genes in the pangenome of *P. ananatis*. Each candidate gene in the accessory genome was scored for its presence and absence. Further, Scoary was used to identify genes that correlated positively with red-onion scale necrosis (indicative of pathogenicity to onion) using the 81 pathogenic- and non-pathogenic *P. ananatis* strains. Scoary predicted 42 genes, including the 14 HiVir/PASVIL cluster genes (*hvaA, pepM, pavC-pavN*) that have been shown to be associated strongly with red-onion scale necrosis or pathogenicity to onion based on stringent *p-*values (S3 Table). A total of 28 genes were identified outside the HiVir/PASVIL cluster. Eight of the 28 genes were annotated and the remaining 20 are hypothetical. Annotated functions of some of the genes included:site-specific tyrosine kinase, N-acetylmuramoyl-L-alanine amidase, TraR/DksA family transcriptional regulator, and HTH-type transcriptional regulator (S3 Table). Out of 14 HiVir/PASVIL cluster genes associated with pathogenicity to onion, three were annotated as hypothetical (*hvaA, pavK* and *pavN*) and the remaining 11 were annotated with functions in metabolite production (Table 2). These 11 genes included *pepM*, coding for phosphonopyruvate mutase; *pavC*, that encodes nitrilotriacetate monooxygenase component A (catalyzes plant-derived aromatic compounds); *pavD* for homocitrate synthase; two genes associated with leucine biosynthesis associated metabolites, including *pavE* for 3-isopropylmalate dehydratase large subunit and *pavF* for 3-isopropylmalate dehydratase small subunit; *pavG*, which codes for SAM dependent methyltransferase; *pavH* for N-acetyltransferase; *pavI* for the ATP-grasp domain containing protein; *pavJ* for MFS transporter; *pavL* which encodes flavin reductase; and *pavM* for carboxylate-amine ligase.

### HiVir/PASVIL and *alt* genes presence and absence

Overall, 80.3% (41 of 51) of the genomes of pathogenic strains of *P. ananatis* contained both HiVir/PASVIL and *alt* conserved clusters. However, none of the non-pathogenic strains contained both of these conserved clusters. Also, none of the pathogenic strains showed the absence of both clusters (Fig 5). The HiVir/PASVIL cluster was conserved in 98% (50 of 51) genomes of the pathogenic strains. The pathogenic strain (PNA_18_9s) showed a partial loss of HiVir/PASVIL genes (*pepM, pavE, pavJ* and *pavK* genes). Absence of these genes could either be due to assembly artifacts or attributed to inconsistent and weak phenotypes (negligible red-scale clearing). Among the 30 non-pathogenic strains, 73.3% (*n* = 22) lacked a complete HiVir/PASVIL cluster and 6.6% (*n* = 2) of the strain genomes showed the presence of only a subset (one or more) of the genes in the HiVir/PASVIL cluster. Interestingly, 20% (*n* = 6) of the non-pathogenic strains possessed a conserved complete HiVir/PASVIL cluster (Fig 5). We also looked into the presence and absence of 11 mega-plasmid borne genes defined as the *alt* cluster. Among the pathogenic strains, the *alt* cluster was conserved in 80.3% (*n* = 41) of the strains, absent in 15.6% (*n* = 8) strains and partially present in two strains (with just one gene present). However, among the non-pathogenic strains, the *alt* cluster was present in 33.3% (*n* = 10) of the genomes, absent in 43.3% (*n* = 13) and partially present (one or two out of 11 *alt* genes present) in 23.3% (*n* = 7) of the strains. Since only one or two genes in the *alt* gene cluster were present in the seven strains (PANS_2_1, PANS_4_2, PANS_99_32, PNA_7_13, PNA_7_14, PNA_98_11, PNA_98_7), these strains were considered negative for the presence of the conserved *alt* cluster (Fig 5). As a result, 66.6% (*n* = 20) of the non-pathogenic strains lacked the *alt* cluster.

**Figure 5.**
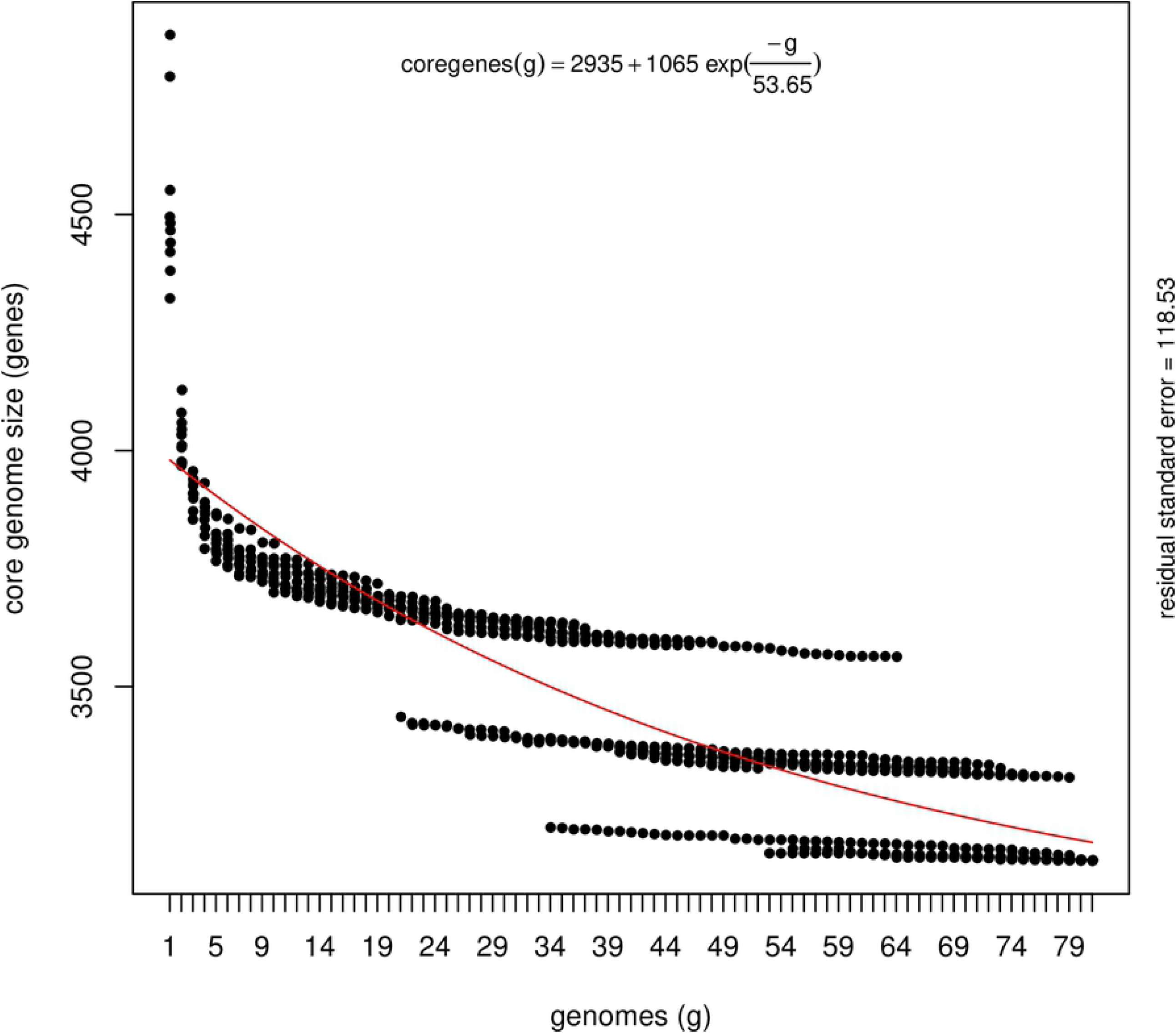
Pictorial representation of presence and absence of HiVir/PASVIL genes associated with red onion scale necrosis and *alt* genes associated with colonization in necrotic zone in pathogenic and non-pathogenic strains of *Pantoea ananatis*.

### Annotation

We annotated core, soft-core, shell and cloud genes that appeared to have roles in biological processes (BP), molecular functions (MF) and cellular components (CC). A total of 2,705 core, 3,293 soft-core, 2,058 shell and 3,503 cloud sequences were annotated successfully, mapped and assigned at least one gene ontology (GO) id and GO slim category (S4, S5, S6, S7 Tables). GO analyses of the top 50 terms revealed that metabolic process represented the most abundant category, followed by cellular processes under BP (Fig 6). Under BP, the cellular amino acid metabolic process was specific to core genes and not present in soft-core, shell and cloud genes (Fig 7). For genes to which MFs could be assigned, catalytic activity was the most abundant category, followed by binding. Transmembrane transporter and oxidoreductase activity, however, was not observed in the shell genes (Fig 7). GO analyses showed that cellular anatomical entities were represented the most abundantly under the CC category (Fig 6 and 7).

**Figure 6.**
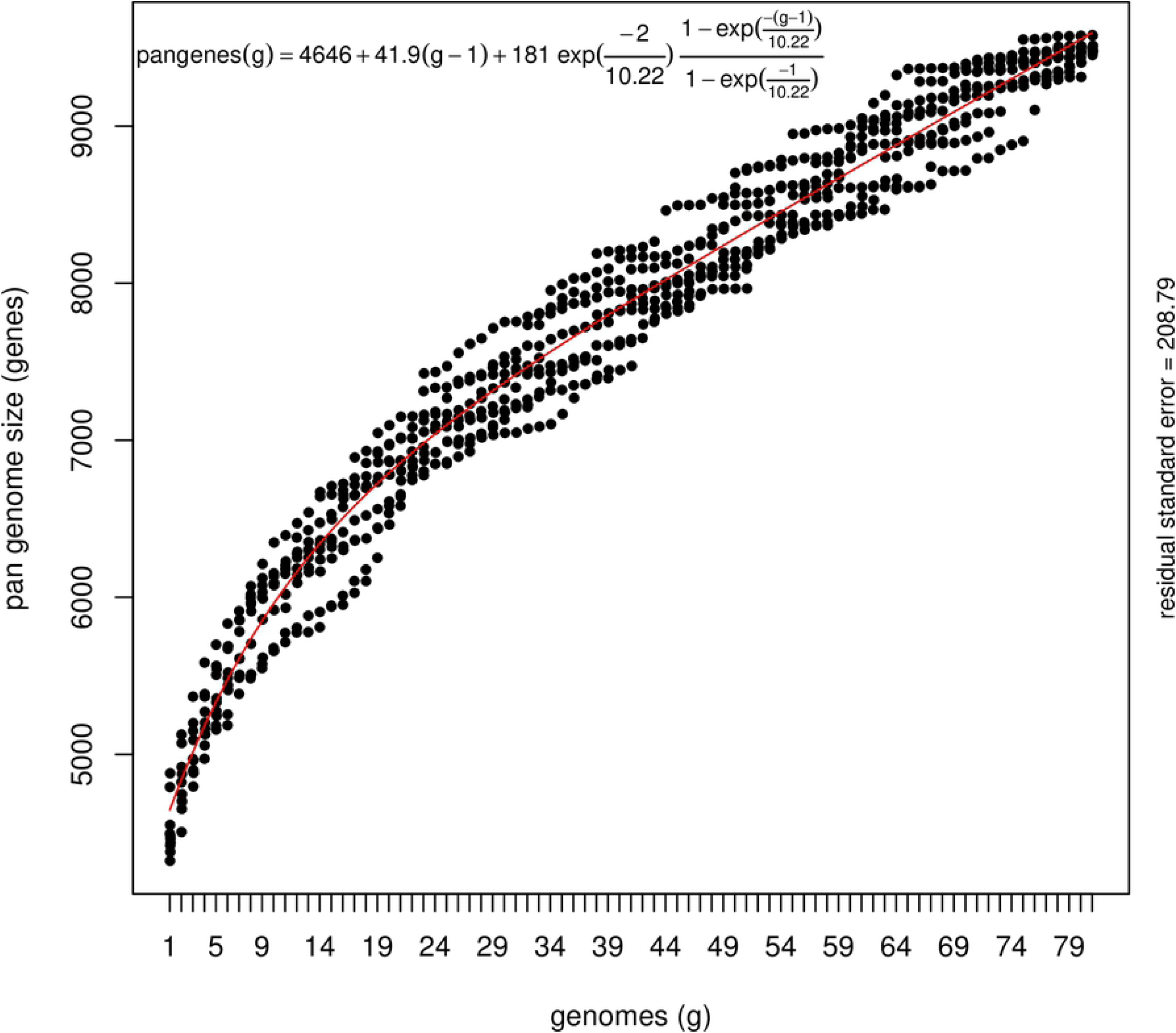
Top 50 GO terms: A bar chart representing the GO terms according to the number of annotated sequences. Panel A-D shows the function of genes assigned to biological process; panel E-H shows the function of genes assigned to molecular function and panel I-L represent the function of genes assigned to cellular component.

**Figure 7.**
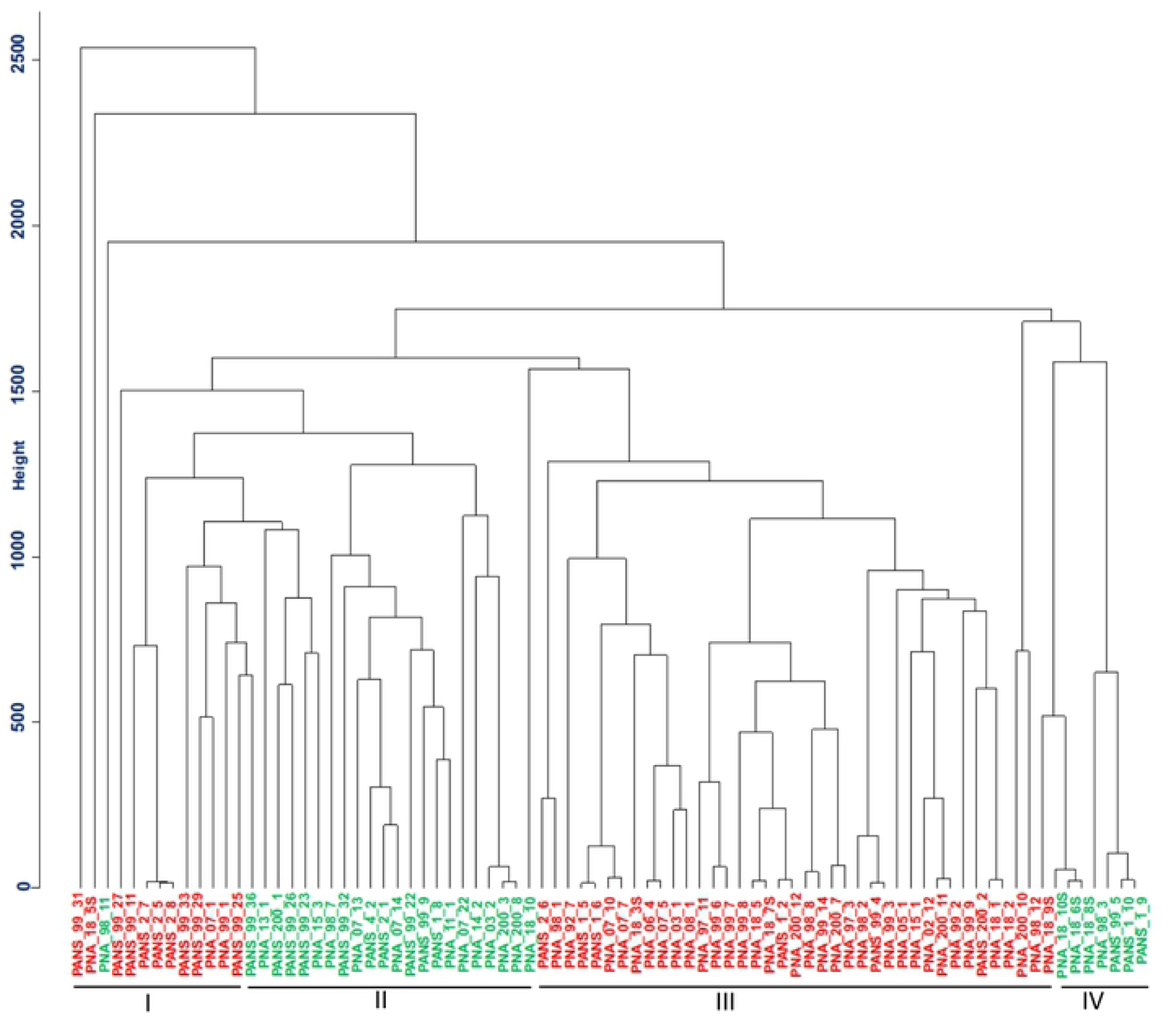
Sequences per GO terms: Annotations of core, soft-core, shell and cloud genes of *Pantoea ananatis* pan-genome. Panel A-D shows the function of genes assigned to biological process; panel E-H shows the function of genes assigned to molecular function and panel I-L represent the function of genes assigned to cellular component.

## Discussion

The complete pan-genome is defined as the total number of non-redundant genes present in a given clade, amounting to a given clade’s entire genomic repertoire, and encodes for all possible ecological niches of the strains examined [37, 38]. A pan-genome typically contains core genes, accessory genes and strain-specific genes. In this study, a core genome represented a conserved set of genes that was present in all of the 81 genomes. The soft core represented genes present in 95% of the total genomes (i.e., ≥76 of the 81 genomes). Defining the soft-core genes aided in obtaining a robust estimate of the core genes. The core and soft-core clusters together represented a pool of highly conserved genes, which could provide insights into the evolutionary history of the *P. ananatis* strains used in this study. The cloud cluster included genes that were strain specific or unique in the pan-genome. Cloud genes, being strain-specific, are unique genes of which only a few genes (1,000 of 6,808) were shared, and only between two genomes. The remaining moderately conserved genes were identified as shell genes. Both the shell and cloud clusters represented the subset of flexible (accessory) genome that reflects life-style, adaptation and evolutionary history of the *P. ananatis* strains to the environments in which they were residing. Earlier pan-genome studies identified 3,875 core genes, 3,876 clusters, 3,750 CDS in seven, eight and 10 *P. ananatis* strains [5, 25, 39]. In this study with a larger set of genomes, we found that the core genome was stabilized with 3,153 gene clusters while the pan-genome expanded continuously as a result of the addition of gene clusters. The openness of a bacterial pan-genome reflects the diversity of the gene pool within the strains of the same bacterial species. The addition of new genomes to an existing pan-genome can significantly alter the pan-genome size of open pan-genomes [40], in contrast to a closed pan-genome.

Another significance of a pan-genome is that it can provide a greater resolution and reconstruct bacterial phylogeny in a more reliable way than single or multiple gene-based phylogeny. The pan-genome provides an overview of the entire gene set (100% of the genomes) of a given population, unlike a 16S rRNA phylogeny that represents only a tiny fraction of the genome (∼0.07%), or multi-locus sequence analysis (MLSA) involving house-keeping genes (∼0.2%). Pan-genome analysis, therefore, provides a greater resolution of phylogeny of the strains than other methods of comparing bacterial strains, as evident in this study. MLSA and rep-PCR assays showed limited genetic diversity, despite high phenotypic variation, among 50 strains of *P. ananatis* [25]. In the same study, Stice et al. [25] demonstrated that PAVs from the pan-genome analysis of 10 strains of *P. ananatis* separated the pathogenic from non-pathogenic strains, which was not observed when the core genome was used. Similar to the study by Stice et al [25], the core and accessory genomes evaluated in this study, when used individually, did not distinguish between pathogenic and non-pathogenic strains; however, the pan-genome PAVs distinguished the pathogenic vs. non-pathogenic strains, except for one non-pathogenic strain that clustered with pathogenic strains, PNA_98_11. In this pan-genome based clustering, we did not find any correlation of the clusters with strains isolated from different years or from different sources (onion vs. other plant hosts) or from different geographic locations, although all the strains originated from the Vidalia region of Georgia. We believe that increasing the number of strains in the pan-genome panel from across onion growing regions and from diverse sources of inoculum may shed some light.

An ANI of ≥95% is a benchmark to classify organisms of the same species [41, 42], whereas genomes of organisms with ANI values of 93-94% suggests the organisms belong to different sub-species [42]. In this study, ANI ranged from 99.0-99.9% and AAI ranged from 96- 99%, which not only suggested low core genome diversity among the 81 *P. ananatis* strains but also confirmed that the strains were all the same species.

Bacterial phenotypes, in general, can be linked to the presence or absence of genes that are inherited through either descent or lateral gene transfer [43]. Previous studies on *P. ananatis* deployed comparative genomics of 2 to 10 strains to identify pathogenicity-related regions in the genome [5, 24, 25, 39]. We conducted pan-GWAS analysis to predict and associate genes related to pathogenicity in onion bulbs. Red-onion scale necrosis has been shown to be an accurate predictor of pathogenicity to onion bulb tissues and, hence, it was utilized as a high-throughput screening to differentiate pathogenic vs. non-pathogenic strains of *P. ananatis*. However, it can be argued that if this assay alone is adequate predictor of pathogenicity both in onion foliage and bulb. We found a strong correlation between bacterial phenotype (pathogenicity) observed in foliar assay vs. red-scale necrosis assay. Hence, this assay was also utilized for the pan-GWAS analysis. We utilized comparative genomics of 81 strains that included 51 onion-pathogenic and 30 strains non-pathogenic to onion. Furthermore, the pan-genome PAVs were utilized to associate onion pathogenicity phenotypes (determined using a red-onion scale necrosis assay) with genes that were associated statistically with pathogenicity to onion. Out of the 42 genes strongly associated with the onion pathogenic phenotype, 14 were identified as a part of HiVir/PASVIL cluster. These genes were annotated as *hvaA, pepM, pavC-N* [23]. These associated genes coded for phosphonate metabolism, metabolism of plant-derived aromatic compounds, monooxygenases, a methyltransferase, leucine biosynthesis and an L-amino acid ligase. Association of HiVir/PASVIL genes using this pan-genome *in-silico* approach corroborated earlier findings of the roles of these genes in *P. ananatis*-pathogenicity on onion [23, 24]. We also using the pan-GWAS on *P. ananatis* to predict 28 novel genes contributing to pathogenicity in onion. Out of the 28 novel genes identified, eight have annotated functions of site-specific tyrosine kinase, N-acetylmuramoyl-L-alanine amidase, TraR/DksA family transcriptional regulator, and HTH-type transcriptional regulator, and the remaining 20 genes are hypothetical. Further functional analysis of these genes will aid in a better understanding of pathogenicity to onion.

Comparative genomic analysis revealed a trend for the presence or absence of complete HiVir/PASVIL cluster genes in pathogenic vs. non-pathogenic *P. ananatis* strains. Among the 51 pathogenic strains evaluated, 50 possessed the complete set of HiVir/PASVIL cluster genes (14 genes; *hvaA*-*pavN*). In the non-pathogenic strains, 73.3% (*n* = 22) of the strains lacked the complete HiVir/PASVIL cluster and 6.6% (*n* = 2) of the strains had only a subset (one or more) of the genes of the HiVir/PASVIL cluster. Interestingly, 20% (*n* = 6) of the non-pathogenic strains possessed a conserved complete HiVir/PASVIL cluster. If the presence of a complete cluster is correlated with pathogenicity to onion, then it is difficult to explain the presence of a complete cluster of genes in these non-pathogenic strains unless the presence of the complete HiVir/PASVIL cluster does not mean the strains are pathogenic on onion. It is possible that the HiVir/PASVIL genes in the cluster in non-pathogenic strains are not expressed or are non-functional, which require further investigation for confirmation.

Phosphoenolpyruvatemutase (*pepM*) is involved in phosphonate biosynthesis [reference?]. Organophosphonates are synthesized as secondary metabolites in certain prokaryotes to function as antibiotics, and can have specialized roles in pathogenesis or signaling [44]. Phosphonate metabolites are derived from phosphonopyruvate, which in turn is formed from phosphoenolpyruvate (PEP) by the action of PEP mutase (PepM). Asselin et al. [24] identified a *pepM* gene as the first pathogenicity factor associated with the fitness of *P. ananatis* as well as with symptom development in infected onion leaves and bulbs. Deletion of *pepM* or inactivation of *pavJ* genes resulted in loss of the ability to cause lesions on onion foliage and yellow onion bulbs. Furthermore, growth of the deletion mutant in onion leaves was significantly reduced compared with the wild-type *P. ananatis* strain. This pan-genome *in-silico* study corroborated the association of *pepM* gene with pathogenicity to onion, using a diverse panel of *P. ananatis* strains from Georgia, USA. The *pepM* gene was present in 50 of 51 pathogenic strains, with the exception strain of PNA 18-9s. This strain also lacked *pavE, pavJ*, and *pavK*. If it is not an assembly artifact, the absence of *pepM* along with four other genes in the HiVir/PASVIL cluster in this strain could be the reason for a compromised red scale necrosis phenotype (weak pathogenicity). This observation also indicated the presence of a potential alternative pathogenicity factor than *pepM*, which requires further investigation. For the non-pathogenic strains of *P. ananatis, pepM* gene was absent in 23 of 30 strains. Despite the presence of *pepM* gene and, in some cases the entire HiVir/PASVIL cluster (six of 30 strains), these strains displayed a non-pathogenic phenotype with the onion red-scale assay. These observations suggest that these genes may be non-functional in these strains, which warrants further research.

A monooxygenase and a flavin reductase enzyme belonging to the two-component non-hemeflavin-diffusible monooxygenase (TC-FDM) family were associated with pathogenicity of *P. ananatis* strains to onion bulb scales in this study The monooxygenase and the reductase associated were nitrilotriacetate monooxygenase coded by *nta*A (similar to *pavC* in the HiVir/PASVIL cluster of *P. ananatis*) and flavin reductase, a flavin:NADH oxidoreductase component of 4-hydroxyphenylacetate (4-HPA) 3-monooxygenase coded by *hpa*C (similar to *pavL* in the HiVir/PASVIL cluster of *P. ananatis*). Nitrilotriacetate monooxygenase was reported previously in the genomic region referred as WHOP (woody host of *Pseudomonas* spp.) in a *Pseudomonas syringae* complex [45]. This region is associated with strains of *P. syringae* that infect woody host plants, and is absent in strains infecting herbaceous tissues. This gene, along with other genes present in the WHOP region, is responsible for the fitness and virulence of *P. savastanoi* pv. *savastanoi* in woody olive trees, but not in non-woody olives [45, 46]. Nitrilotriacetate monooxygenase is known to catabolize plant-derived aromatic compounds and help bacteria to adapt to woody host tissues [47]. On the contrary, center rot of onion is caused by *P. ananatis* moving into onion bulbs from infected foliage or neck tissue; therefore, it was intriguing to find this gene associated with pathogenicity in onion, an herbaceous plant. The 4-hydroxyphenylacetate (4-HPA) 3-monooxygenase reductase uses NAD(P)H to catalyze the reduction of free flavins that diffuse to the oxygenase component for oxidation of the substrate (aromatic or non-aromatic compounds) by molecular oxygen.

The HiVir/PASVIL gene *pavG* in *P. ananatis* has an annotated function for a class-I S-adenosyl-L-methionine (SAM)-dependent methyltransferase. We presume that *pavG* is responsible for the esterification of phosphonates synthesized in *P. ananatis*, led by *pepM*, based on the fact that the di-anionic form of phosphonates interferes with the metabolic intermediates and carboxylates of antibacterial compounds [48-50]. To counteract this problem, microbes either synthesize phosphinites (with a double bond between the C and P instead of a single bond) or carry out esterification of phosphonates. Phosphonate esterification appears to be an obvious mechanism operational in *P. ananatis* because of the presence of *pavG* in the HiVir/PASVIL cluster. SAM dependent O-methyltransferase has been shown to methylate a variety of phosphonates (1-hydroxyethylphosphonate, 1,2-dihydroxyethylphosphonate, and acetyl-1-aminoethylphosphonate) [49]. Therefore, there is a high possibility of involvement of SAM methyltransferase in methylation of the phosphonate produced in *P. ananatis*. Further studies are required to characterize the type of phosphonate and its methylation in order to understand the mechanism of SAM methyltransferase and *pepM* in causing red-scale necrosis. Another role that *pavG* could be playing is methylation of other HiVir/PASVIL genes that renders them inactive despite their presence in the cluster. We hypothesize that the inactivity of HiVir/PASVIL genes may be due to methylation of genes carried out by SAM dependent methyltransferase in non-pathogenic strains of *P. ananatis*, implying a secondary role of *pavG* in strains non-pathogenic to onion. Methylation profiling will help evaluate this hypothesis.

The HiVir/PASVIL gene *pavI* is similar to *RizA*, an L-amino-acid ligase (LAL) from *Bacillus subtilis* that participates in the biosynthesis of rhizocticin, a phosphonate oligopeptide antibiotic and possess L-arginyl-L-2-amino-5-phosphono-3-cis-pentenoic acid [51]. Although, the functional role of *pavI* is yet to be characterized in *P. ananatis*, it may play a role in the formation of anti-microbial secondary metabolites of ‘phosphonate derivatives’. LAL is a member of the ATP-dependent carboxylate–amine/thiol ligase superfamily [52], and catalyzes the ligation reaction which involves an aminoacyl-phosphate intermediate, in an ATP-dependent manner [53]. LALs contain the ATP-grasp fold, which is composed of three conserved domains referred to as the A-domain (N-terminal domain), the B-domain (central domain) and the C-domain (C-terminal domain). These three domains commonly grasp the ATP molecule, and also provide binding sites for the Mg2+ ion and the amino-acid substrate [54].

The pan-GWAS approach used in this study did not associate the *alt* cluster of genes with the onion pathogenic phenotype using the red-scale necrosis assay. This may be because the phenotyping was based only on the necrotic scale clearing assay thathas been shown to be induced by the HiVir/PASVIL cluster [reference]. However, the *alt* cluster comes into play after the onset of necrosis, when endogenous antimicrobial sulfur compounds are produced by damaged onion cells. In this scenario, the *alt* cluster helps *P. ananatis* survive and colonize onion plants. *P. ananatis* uses 11 *alt* cluster genes associated with sulfur metabolism to impart tolerance to the thiosulfinate ‘allicin that is produced by damaged onion cells [26]. The presence of the *alt* cluster in 80% (*n* = 41) of the onion pathogenic strains, and its absence in 67% (*n* = 20) of the non-pathogenic strains, suggests some potential involvement in bacterial virulence. However, the *alt* cluster alone is not responsible for the onion pathogenic phenotype, as 33% (*n* = 10) of the non-pathogenic strains did not exhibit any evidence of pathogenicity in the onion red scale assay despite possessing a completely conserved *alt* cluster.

In this study, we demonstrated the use of a pan-GWAS approach to predict genes associated with onion-pathogenicity in *P. ananatis*. The pan-GWAS predicted 42 genes, including 14 genes from the previously reported HiVir/PASVIL cluster that is associated with pathogenicity, and 28 novel genes that were not previously known to be associated with pathogenicity of *P. ananatis* to onion. Results of the pan-genome analysis led to the conclusion that HiVir/PASVIL genes are associated with onion-pathogenicity based on the red-scale necrosis assay, and the *alt* gene cluster alone is not responsible for pathogenicity to onion. Also, HiVir/PASVIL gene expression is potentially regulated, and the mere presence of the HiVir/PASVIL cluster does not guarantee a strain is pathogenic to onion. In addition, a large repertoire of accessory genes identified in these strains may aid *P. ananatis* in diverse niche-adaptation and potentially in host-range expansion. The pan-GWAS pipeline can be deployed to characterize *P. ananatis* strains pathogenic to other plant hosts.

## Materials and methods

### Bacterial strains, identification and culture preparation

Eighty-one *P. ananatis* strains used in this study were isolated from onion foliage, bulb and seeds as well as from weeds and thrips throughout the state of Georgia (Table 1). These strains were stored in a 15% aqueous glycerol solution at -80°C. The source, year of isolation, and county of origin in Georgia for these strains are listed (Table 1). Strains were identified as *P. ananatis* by their colony morphology and physiological characteristics as Gram negative, facultatively anaerobic, positive for indole production, negative for nitrate reductase and phenylalanine deaminase and by PCR amplification of a 398 base pair (bp) fragment using *P. ananatis*-specific primers (Walcott et al., 2002). Strains that were isolated from onion plants were designated as ‘PNA’ and strains from non-onion sources (e.g., weeds or thrips) were identified as ‘PANS’.

Inoculum was prepared by transferring single colonies of each bacterial strain from 48-h-old cultures on nutrient agar (NA) medium to nutrient broth (NB). The broth was shaken overnight on a rotary shaker (Thermo Scientific, Gainesville, FL) at 180 rpm. After 12 h of incubation, 3 ml of each bacterial suspension were centrifuged at 6,000 × *g* (Eppendorf, Westbury, NY) for 2 mins. The supernatant was discarded and the pellet was re-suspended in sterile water. Inoculum concentration was adjusted using a spectrophotometer (Eppendorf, Westbury, NY) to an optical density of 0.3 at 600 nm [≈1 × 10^8^ colony forming unit (CFU)/ml]. The bacterial suspension was diluted serially in sterile distilled water to obtain the desired concentration.

### Phenotyping of *P. ananatis*: red onion scale necrosis assay as a measure of pathogenicity to onion

The red onion scale necrosis assay was conducted as described by Stice *et al*. [25]. Briefly, red onion bulbs (cv. Red Creole) were each surface-disinfested by removing the outer, dry scales and then spraying the outer exposed fleshy scales with a 3% sodium hypochlorite solution using a spray bottle. This was followed by blotting the outer scales dry with a sterile paper towel. Later, the bulbs were sprayed with sterile distilled water and blotted again with a sterile paper towel. Surface-disinfested onion scales were each sliced into approximately 6 cm × 3 cm (length × width) segments using a surface-disinfested knife. Each onion scale piece was pierced on the inner surface with a pipette tip, and 10 μl of the relevant bacterial suspension (1 × 10^6^ CFU/ml) was deposited on the wounded tissue. Scales were maintained in petri plates containing autoclaved paper towels moistened with sterile water. Petri plates were kept in an aluminum tray covered with a plastic lid. The inoculated onion scales were incubated at room temperature for 96 h in the dark, after which the size of the necrotic, pigment-clearing zone was recorded as a measure of pathogenicity. Strains that cleared the red anthocyanin pigment and caused necrosis were classified as pathogenic, and those that did not were classified as non-pathogenic to onion. Three replicate onion scale pieces were used to test each strain, and the experiment was repeated twice (a total three experiments). Onion scales inoculated with sterile water and a known pathogenic strain of *P. ananatis* (PNA 97- 1) served as negative and positive control treatments, respectively.

To confirm that symptoms on the onion scale were caused by *P. ananatis*, bacteria from symptomatic scale tissue (*n* = 3) were isolated from the margins of the necrotic area and healthy tissue, and streaked onto Tryptic soy broth agar (TSBA) and incubated for 48 h at 28°C. Yellow-pigmented colonies were selected to test for genus and species identity using physiological tests and a species-specific TaqMan-based polymerase-chain reaction (PCR) assay [13] for *P. ananatis*. Briefly, presumptive *P. ananatis* colonies were picked using a sterile inoculation loop and suspended separately in 2 ml micro-centrifuge tubes, each containing 25 μl of sterile deionized water. The bacterial suspension was heated (Modular Dry Block Heaters, Cole Parmer, IL) for 3 min at 95°C. A suspension (5 µl) was amplified in 20 μl of PCR master-mix containing 10 mMmTris-HCl (pH 9.0), 50 mM KCl, 0.1% Triton X-100, 1.5 mM MgCl_2_, and 0.2 mM of each nucleotide (dATP, dCTP, dGTP, and dTTP), 25 μM each of primer PanITS1 (5′-GTCTGATAGAAAGATAAAGAC-3′) and EC5 (5′-CGGTGGATGCCCTGGCA-3′) and 10 μM of TaqMan probe 6-FAM TAGCGGTTAGGACTCCGCCCTTTCA-BHQ. The PCR reaction was conducted in a Cepheid Smart Cycler (Sunnyvale, CA) using the following thermal profile: denaturation at 95°C for 180 s, 35 cycles each of denaturation at 95°C for 15 s, and annealing at 60°C for 40 s. Samples with cycle threshold (Ct) values <35 were considered positive for *P. ananatis*.

### DNA isolation and library preparation for whole genome sequencing of *P. ananatis*

*P. ananatis* strains (*n* = 81) were streaked individually onto NA and incubated for 48-h at 28°C. After incubation, a single colony of each strain was picked and placed into 3 ml of Luria Bertani broth. The resulting broth was shaken overnight on a rotatory shaker (Thermo Scientific, Gainesville, FL) at 180 rpm at 30°C. After incubation, broth (1.5 ml) cultures were each centrifuged in a microcentrifuge tube (2 ml) at 6,000 × g. The supernatant was discarded and the bacterial pellet suspended in 1 ml of sterile water, from which DNA was isolated. DNA isolation was done using a DNeasy ultra clean microbial DNA extraction kit (Qiagen, Germany) using the manufacturer’s instructions. DNA samples were quantified (ng/µl) using a Nanodrop (Thermo Scientific, Gainesville, FL).

For Illumina Nextera library preparation, a total of 100 ng DNA from each bacterial strain was used according to KAPA Hyper Prep kit (KAPA Biosystems, MA, USA) at the Georgia Genomics and Bioinformatics Core (GGBC), University of Georgia, Athens, GA, USA. Briefly, the genomic DNA from each bacterial strain was fragmented followed by end repairing and A-tailing, which produced end-repaired 5’-phosphorylated, 3’-dA-tailed DNA fragments. Adapters were ligated resulting in a 3’-dTMP overhang ligated to 3’-dA-tailed molecules. Post-ligation cleanup was performed to remove unligated adapter and/or adapter-dimer molecules from the library before library amplification. Library amplification was done employing a high-fidelity, low-bias PCR assay to amplify library fragments with appropriate adapters on the ends. Dual indexing was done during the library preparation and PCR amplification. Dual indexing (by introducing indexes into both library adapters) was done to overcome the occurrence of mixed clusters on the flow as this is a predominant source of error while multiplex sequencing. Libraries for the 81 bacterial strains were pooled and sequenced on the Illumina Nextseq500 using a high output run. All samples were sequenced to produce 150 bp paired end reads.

### Read data filtering

FastQC was run to assess the raw fastq files. Overall, data quality was good with typical drop off in quality at the ends of the reads. The number of raw reads ranged from 8.4 million to 18.2 million PE reads. The read data were filtered to remove low quality reads/bases and trimmed for reads containing primer/adaptor sequences using Trimmomatic (v 0.36) in paired end mode [55]. In addition, all 5’ and 3’ stretches of ambiguous ‘N’ nucleotides were clipped. Reads were trimmed twice to ensure adequate quality of data before downstream analyses. To ensure high quality reads for downstream analysis, a second trimming step was employed. Trimmed data were re-assessed using FastQC. Second trimmed data loss averaged 2.7%. These data were used for genome assembly followed by pan-genome analyses.

### Genome assembly and pan-genome analyses

Trim2 reads were assembled using the SPAdes (v 3.11.1) assembler [56]. Both the paired and unpaired data were used in assembly at default settings. Assembly files were submitted to NCBI under the bioproject identity PRJNA624643. The assembled contigs were also submitted to LINbase to obtain life identification numbers (LIN) (Table S8). The scaffolds of the respective 81 *P. ananatis* strains were annotated using prokka (v 1.13) [57] to produce gff3 (general feature format) and gbk (gene bank format) files. The gbk files were used for pan-genome analyses by using get_homologues [58]. These gbk files were used to get the syntenic sequence clusters by get_homologues.pl using OrthoMCL (OMCL) algorithm. The syntenic clusters generated were used to a develop a pan-genome matrix showing presence and absence variants (PAVs) using compare_clusters.pl, and the pan-genome matrix was used to classify the genes into core, soft-core, shell and cloud genes using parse_pangenome_matrix.pl (auxiliary script of get_homologues.pl). Core genes were defined as those present in all 81 genomes and accessory genes were present in a subset of the 81 strains. The accessory gene cluster was further divided into soft-core, shell and cloud gene clusters. Soft-core genes occurred in 95% of the genomes. Cloud genes were present in ≤2 genomes and shell genes comprised of remaining genes [58]. Distribution of cluster sizes as a function of the number of genomes these clusters contained was displayed using R with parse_pangenome_matrix.pl. Gower’s distance matrix was generated using the tab delimited pan-genome PAV file as input by executing the shell script hcluster_pangenome_matrix.sh (auxiliary script of get_homologues) when used to call R function hclust. The presence and absence of 14 HiVir/PASVIL and 11 *alt* cluster genes were determined by looking at the cluster of genes using compare_clusters.pl program and blastn search respectively. HiVir/PASVIL genes that were clustered from different strains were considered present in those strains. Each of the *alt* cluster sequence was subject to blastn against each of the 81 genome assemblies. If a blast hit was found against a genome, that gene was considered present. The *alt* cluster was considered conserved in a given strain if all 11 genes were present, and was considered absent if anywhere from 9 to 11 genes were absent. Similarly, if all 14 genes of the HiVir/PASVIL cluster were present in a strain, then it was considered conserved, otherwise it was rated as absent if all some or all of the genes were absent in a particular strain. The presence and absence of these genes was plotted as heat maps.

### Average nucleotide/amino acid identity, pan-genome wide association and annotation

The Get_homologues perl program was used to estimate the average nucleotide (ANI) and amino acid (AAI) identities of CDSs and the proteins coded by the CDSs among all individual strains of the pan-genome. The resulting ANI (-a ‘CDS’) and AAI pan-genome matrices obtained using get_homologues.pl were used to plot the heat maps with the shell script plot_matrix_heatmap.sh. For AAI, BLASTP scores were calculated among protein sequences. For association analysis, the pan-genome matrix was used with the phenotyping data using Scoary, a python program [43]. Scoary was used to calculate associations among genes in the pan-genome and the red-scale necrosis assay (a qualitative; pathogenic vs. non-pathogenic association). The output of this program comprised of a list of genes sorted by strength of association with these traits. Genes with a naïve p-value <0.05, a Bonferroni p-value <0.05 and corrected p-value (Benjamini-Hochberg) of association <0.05 were considered significant. The predicted core, soft-core, shell and cloud gene sequences were annotated by doing a blastx search (E value ≤1e−5) against the NR database. The blast output was generated in an XML format that was used as input for Blast2GO to assign gene ontology (GO) terms and assignment of KEGG pathways [59].

## Acknowledgments

We thank Matthew Tyler Garrick, Medora Hoopes, Walt Sanders and David Abgott for technical assistance in the field and laboratory.

## Conflict of interest

The authors declare that they have no conflict of interests.

## Authors’ contribution

GA and BD conceived the project, GA performed the bioinformatics analyses, and complied the manuscript. DC maintained the bacterial cultures, isolation of strains and phenotyping of the 81 strains. SPC and BHK contributed in planning and designing the experiment and manuscript revision. BKM and SNV contributed to the discussion. RG and BD designed and finalized the manuscript. BD planned the project, secured extramural funds, and revised and submitted manuscript.

## Data availability statement

Assemblies with over 200 bp contig size have been deposited to the NCBI (PRJNA624643).

## Funding

This study was supported in part by resources and technical expertise from the Georgia Advanced Computing Resource Center, a partnership between the University of Georgia Office of the Vice President for Research and Office of the Vice President for Information Technology. This work is partially supported by the Specialty Crop Block Grant AWD00009682. This work is also partially supported by the Specialty Crops Research Initiative Award 2019-51181-30013 from the USDA National Institute of Food and Agriculture. Any opinions, findings, conclusions, or recommendations expressed in this publication are those of the author(s) and do not necessarily reflect the view of the U.S. Department of Agriculture. The University of Georgia is an equal opportunity provider and employer.

## Supporting information

**S1 Fig.**
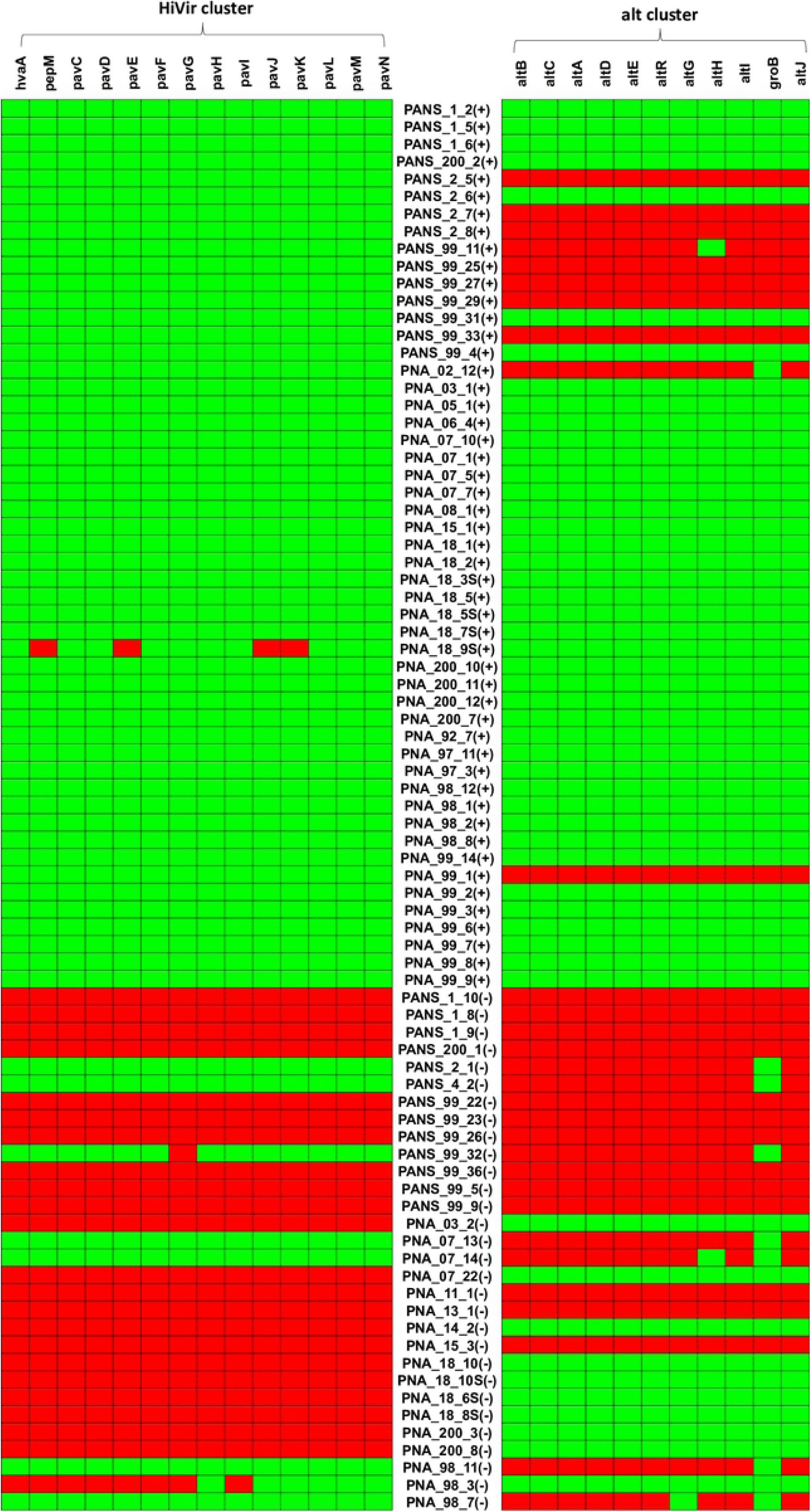
Raw-read data generated, and filtered reads retained after stringent quality filtering using Trimmomatic.

**S2 Fig.**
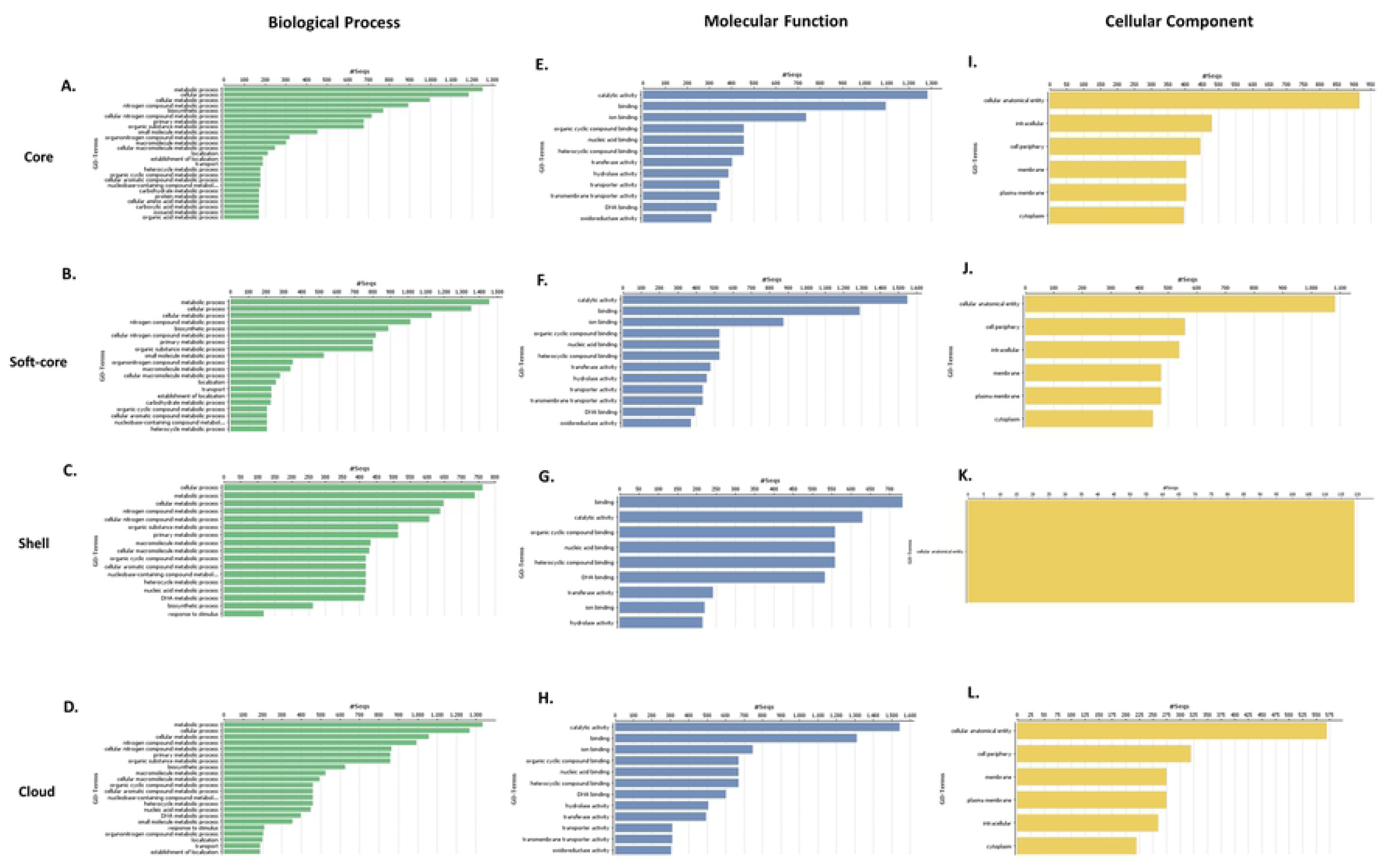
Dendrogram of 51 red-onion scale necrotic and 30 red-onion scale non-necrotic strains of *P. ananatis* based on the conserved core gene set. Strains highlighted in green are non-pathogenic and ones in red are pathogenic.

**S3 Fig.**
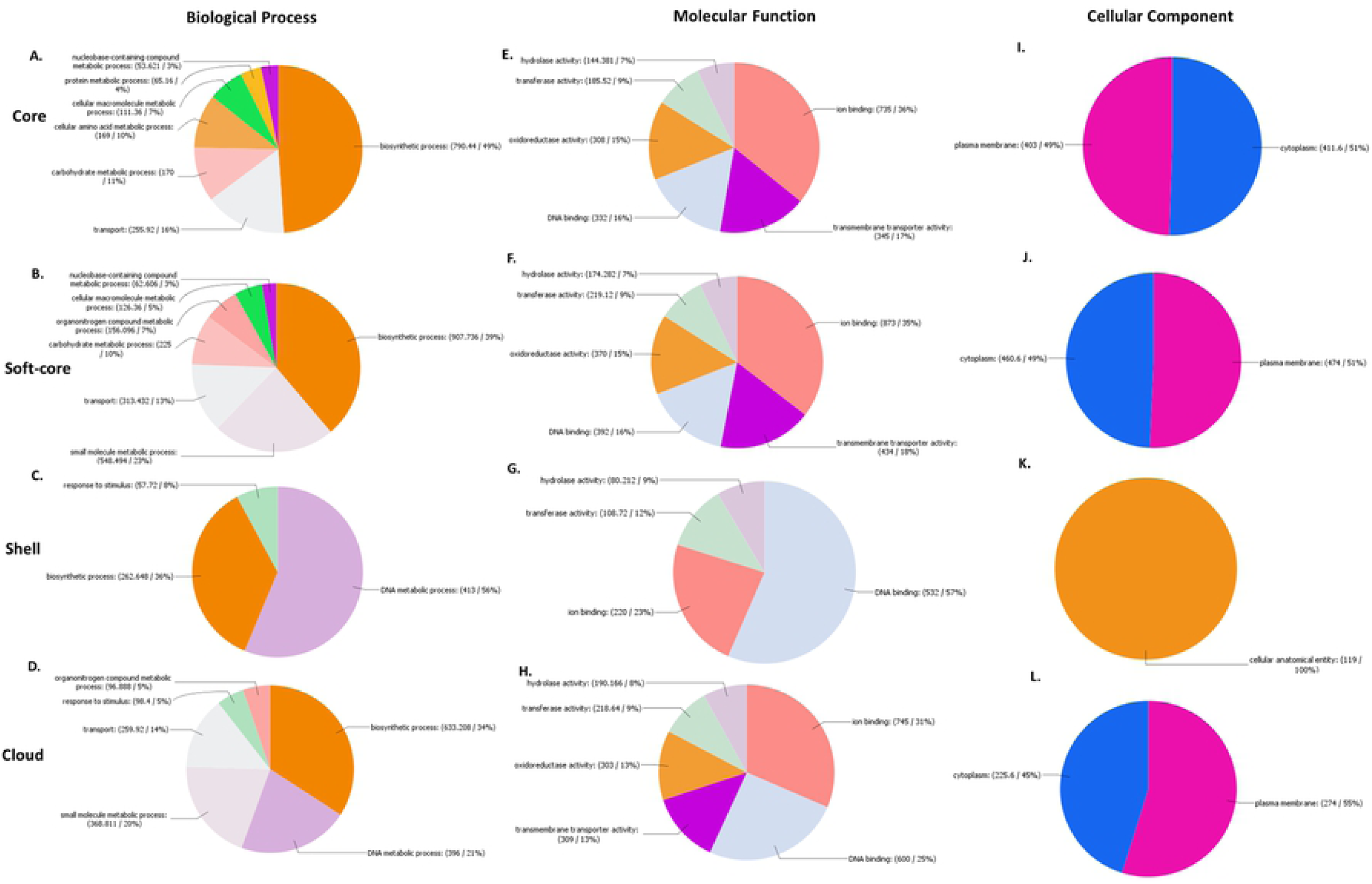
Dendrogram of 51 red-onion scale necrotic and 30 red-onion scale non-necrotic strains of *P. ananatis* based on shell genes. Strains highlighted in green are non-pathogenic and ones in red are pathogenic.

**S4 Fig.**
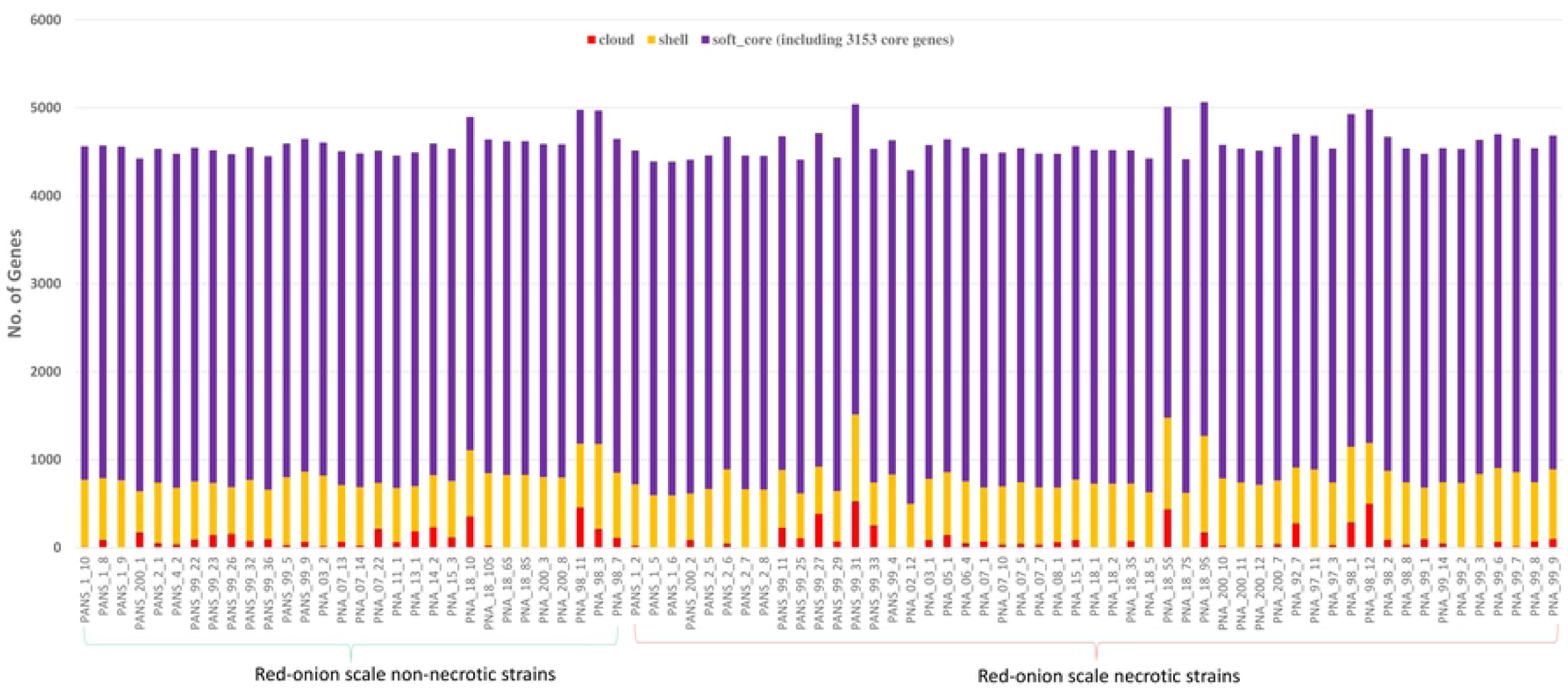
Dendrogram of 51 red-onion scale necrotic and 30 red-onion scale non-necrotic strains of *P. ananatis* based on cloud genes. Strains highlighted in green are non-pathogenic and ones in red are pathogenic.

**S1 Table**. Details of raw and filtered data generated for each sample of *P. ananatis* used in this study.

**S2 Table**. Total number of core and accessory genes shared by each strain of *P. ananatis* used in this study.

**S3 Table**. List of 42 genes associated with red-onion scale necrosis caused by pathogenic strains of *P. ananatis* based on *P*-values.

**S4 Table**. Annotation and mapping of representative core genes.

**S5 Table**. Annotation and mapping of representative soft-core genes.

**S6 Table**. Annotation and mapping of representative shell genes.

**S7 Table**. Annotation and mapping of representative cloud genes.

**S8 Table**. Life Identification Number (LIN) Ids assigned to *P. ananatis* strains submitted to LIN base.

